# The rhizosphere of *Picea abies* is a hotspot of terpenoid production

**DOI:** 10.64898/2026.08.12.744374

**Authors:** Mirjam Meischner, Alexej Steuerle, Riikka Rinnan, Christiane Werner

## Abstract

Forest soils are an important source of volatile organic compounds (VOCs), yet little is known about how different tree species influence soil VOC emissions and the role of rhizosphere processes in mediating VOC release form roots.

We analysed soil VOC emissions from the soil surface and bulk soil as well as from roots with intact rhizosphere and washed roots of *Picea abies* and *Fagus sylvatica*. Tree saplings were grown on natural forest soil, and VOC emissions and gas exchange of soils and roots were measured under controlled conditions using online gas analysers integrated into an automated system. To assess the contribution of rhizosphere soil and microbial communities to root VOC emissions, roots were analysed (a) without washing, preserving the rhizosphere, (b) water-washed, and (c) ethanol-washed (70 vol%) to minimize microbial contributions.

Species-specific VOC emission patterns were observed in both soils and roots. *P. abies* showed higher total emission rates and a more diverse, terpenoid-rich VOC profile dominated by α-pinene, β-pinene, β-myrcene, and α-phellandrene than *F. sylvatica*. Notably, these differences were evident not only at the soil surface but also in root and litter free bulk soil. Root washing further revealed that the rhizosphere is a hotspot of terpenoid production in *P. abies*, with significantly higher monoterpenoid emissions from unwashed roots than from water or ethanol-washed roots.

This study demonstrates how tree species shape net soil VOC emissions, potentially leading to cascading effects on atmospheric VOC concentrations, and highlights the importance of the rhizosphere in regulating belowground VOC production.

## Introduction

Biogenic volatile organic compounds (VOCs) are pivotal to plants for protection against biotic and abiotic stressors (Kesselmeier & Staudt, 1999; Holopainen, 2004; Loreto *et al*., 2014). These small, lipophilic molecules are characterized by a high vapor pressure (Pichersky *et al*., 2006; Dudareva *et al*., 2006), and their composition and concentration are dependent on the stress status of the plant (Karban & Myers, 1989; Niinemets, 2010; Werner *et al*., 2021), varying significantly across species (Owen *et al*., 2001; Haberstroh *et al*., 2018) and plant organs (Bertoli *et al*., 2004; Daber *et al*., 2025). VOC emissions by aboveground plant organs, i.e. flowers (Knudsen *et al*., 2006; Schiestl, 2010) and leaves (Kesselmeier & Staudt, 1999; Tholl *et al*., 2006; Niinemets *et al*., 2011), are well documented and fulfill important functions such as attracting or repelling insects (Karban & Myers, 1989; Erb & Reymond, 2019) or reducing cell damage (Singsaas *et al*., 1997; Velikova *et al*., 2012), among other functions. However, the ecology and production of volatiles released by plants into the soil remains poorly understood (Peñuelas *et al*., 2014; Delory *et al*., 2016). This represents a significant gap in our understanding of plant VOCs, given that the root system accounts for 10-30% of the total plant biomass in temperate forests (Bolte *et al*., 2004; Mokany *et al*., 2006), and even exceeds the aboveground biomass by a factor of four on average in grasslands and arid ecosystems (Mokany *et al*., 2006). Furthermore, it has been shown, that plants release 5–21% of their assimilates into the rhizosphere (Jones *et al*., 2009). The rhizosphere is defined as the interface between roots and soil and is characterized by a high microbial activity (Bakker *et al*., 2013) and numerous biotic interactions mediated by a diverse array of volatile and non-volatile organic compounds (Junker & Tholl, 2013; Massalha *et al*., 2017; Mathieu *et al*., 2024). However, due to methodological constraints, volatile compounds often receive little attention in studies on root exudates (Baetz & Martinoia, 2014; Oburger & Jones, 2018; Wang *et al*., 2021).

Early research on root VOCs relied on extractions of essential oils from dried or frozen roots (Ryan & Guerin, 1982; Rohloff, 2002). Rohloff (2002) revealed a high chemical diversity and abundance of VOCs in *Rhodiola rosea* roots, including terpenoids, alcohols, aldehydes and acids. While the quantification of stored VOCs did not provide a direct measure of actual emission rates, it led to further studies that established the ecological importance of root VOCs as signaling molecules based on their characteristic volatility. For instance, it was shown through olfactometer experiments that herbivorous nematodes are specifically attracted to insect-damaged roots (Rasmann *et al*., 2005; Ali *et al*., 2010). (E)-β-caryophyllene, gijerene and pregeijerene were identified as the terpenoids mediating the interaction between damaged roots and entomopathogenic nematodes, highlighting the critical role that root VOCs play in belowground biotic interactions (Rasmann *et al*., 2005; Ali *et al*., 2010). With the advent of novel measurement techniques, high-resolution time series using proton-transfer-reaction time-of-flight mass spectrometry (PTR-TOF-MS) have revealed pronounced time-dependent changes in root VOC emissions following root damage and herbivory (Danner *et al*., 2012; Crespo *et al*., 2012), as well as a significant effect of aboveground stress on root VOC production and emission rates (Steeghs *et al*., 2004; Meischner *et al*., 2026). So far, however, root VOC research has focused heavily on herbaceous species, such as *Brassica spp.* (Danner *et al*., 2012; Crespo *et al*., 2012; Voyard *et al*., 2024) and *Solanum spp.* (Gulati *et al*., 2020; Voyard *et al*., 2024), leaving a significant gap in our knowledge of root VOC emissions from woody species.

Experimental plants for studies on root VOCs are mostly grown in soil-free laboratory conditions (Steeghs *et al*., 2004) or sand-rich artificial soils (Crespo *et al*., 2012) and therefore may not accurately reflect the complex interactions between roots and their microbiome in a natural soil. An alternative approach involves non-invasive measurement of VOCs from the soil surface using the open-bottom chamber method (Aaltonen *et al*., 2013; Gray *et al*., 2014; Mäki *et al*., 2019a; Pugliese *et al*., 2022; Mu *et al*., 2022; Rinnan, 2024; Kreuzwieser *et al*., 2025). VOC emissions measured in this way represent the VOCs produced by plants and the soil microbiome minus VOCs that are dissolved in soil water, attached to soil particles, or, importantly, have been degraded by soil microorganisms (Insam & Seewald, 2010; Tang *et al*., 2019). While measurements of net soil VOC emissions provide valuable information on the contribution of soils to the ecosystem-atmosphere exchange of VOCs, the source of emitted VOCs cannot be identified. Especially the degradation of VOCs by soil microorganisms might considerably reduce the amount of root VOCs that are released by soils (Albers *et al*., 2018; Jiao *et al*., 2023), with the sink capacity of soils being strongly dependent on the soil humidity and temperature (Pugliese *et al*., 2022). Accordingly, Voyard *et al*. (2024) demonstrated that root VOC emissions from *Brassica napus* L. and *Solanum lycopersicum* L. increased drastically upon the removal of soil. However, the influence of the rhizosphere on these emissions, i.e. whether the rhizosphere acts as a net source or sink of root-emitted VOCs remains an open question. Resolving this issue is critical for advancing our understanding of the processes at the root-soil interface.

The objective of this study was to investigate the processes driving soil and root VOC emissions in two woody plant species, focusing specifically on the role of the rhizosphere. Although the rhizosphere represents a small volume compared to the bulk soil, its high microbial activity is expected to exert a considerable influence on net root VOC emissions. We studied two widely distributed and economically significant tree species in Central Europe: *Picea abies*, a conifer characterized by high VOC emission rates and substantial VOC storage pools (Fäldt *et al*., 2003; Filella *et al*., 2007), and *Fagus sylvatica*, a broadleaved species with relatively low VOC emission rates and labile storage pools (Dindorf *et al*., 2006; Holzke *et al*., 2006). These contrasting aboveground VOC emission profiles are also reflected in the root emissions, as demonstrated in a previous study (Meischner *et al*., 2025). Here we test the hypotheses that (i) species-specific differences in root VOCs will shape net soil VOC emissions but will not persist in litter and root free soil and that (ii) in both species the rhizosphere will act as a net sink for VOCs due to microbial degradation (Asensio *et al*., 2007), such that washed roots will exhibit overall higher emission rates than roots with an intact rhizosphere. To test these hypotheses, VOC emissions of *F. sylvatica* and *P. abies* were measured from the soil surface, excavated roots, and root and litter-free bulk soil under controlled conditions. In order to investigate the influence of the rhizosphere, three treatments were applied: a) no washing, to maintain intact rhizosphere soil and root adhering microbial communities b) water washing, to remove rhizosphere soil and c) ethanol washing (70 vol. % ethanol), to minimize microbial abundance on the root.

## Material and methods

### Plant and soil material

In February 2023, 24 saplings of *Fagus sylvatica* L. and *Picea abies* (L.) H. Karst., along with forest soil, were collected from a forest dominated by these two species near Ettenheimmünster in the Black Forest, South-West Germany (N48.2562°, E7.9220°). The soil at the collection site is classified as Cambisol, developed from silty and loamy soils over red sandstone parent material (LGRB, 2021). The soil texture is a silty-loam to clay type with a skeleton content of approximately 20% in the upper soil layer (Kinzinger *et al*., 2024). The plants were potted into their original forest soil to preserve the natural root microbiome. Plants were gardened outdoors at the University of Freiburg (N48.01418°, E7.8336°) until they were transferred to walk-in climate chambers (Thermotec, Weilburg, Germany) for acclimatization in August 2025. By this time, *F. sylvatica* and *P. abies* had a mean height of 73.5 ± 5.0 cm (mean ± SD) and 63.3 ± 4.9 cm and a mean root crown diameter of 17.0 ± 1.1 cm and 19.5 ± 2.2 cm, respectively. The climate chamber conditions were set to a day length of 12 hours (with an additional 30 minutes of dusk and dawn), a light intensity of 700 µmol m^-2^ s^-1^ photosynthetically active radiation active radiation at mid-shoot height, a relative air humidity of 50 % and day/night air temperatures of 25°/15°C.

### Experimental procedure and treatments

All experimental plants and their associated soils underwent four distinct measurement cycles. These began with VOC analyses of the soil surface, followed by two sequential root measurements (T1 and T2). To evaluate the impact of the rhizosphere soil, plants were randomly assigned to either a ‘rhizosphere’ group or an ‘ethanol’ group. In the ‘rhizosphere’ group, roots were measured first with the rhizosphere soil attached and subsequently washed with water and then measured again. In the ‘ethanol’ group, roots were first washed with water and then treated with ethanol to further reduce the amount of root adhering microbes (Fig. 1). This allowed comparisons between roots with an intact rhizosphere and water-washed roots (T1), as well as between ethanol-washed and water-washed roots (T2). As a final step, the VOC emissions from the root free bulk soil were analyzed (Fig 1). In total, the experiment includes the three root treatments ‘rhizosphere’, ‘water-washed’, and ‘ethanol-washed’, as well as the soil treatments ‘bulk soil’ and ‘soil surface’ (Fig 1).

**Figure 1:**
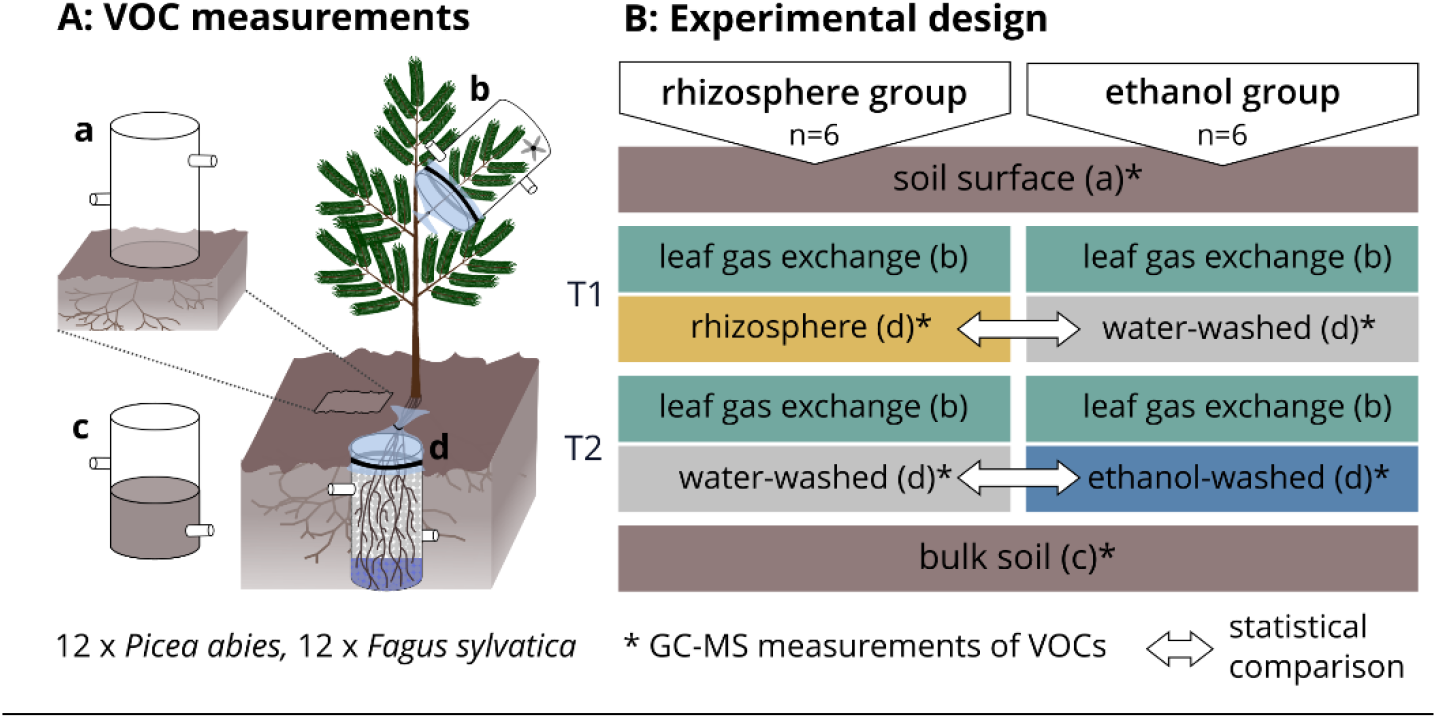
Soil, root and shoot VOC emissions and gas exchange (CO_2_ and H_2_O) were measured continuously with plant or soil enclosures (b, c, d) or open bottom chambers (a) connected to an automated flow-through gas exchange measurement system (left panel). Measurements were performed in four cycles for each group (rhizosphere, ethanol) and species (*F. sylvatica*, *P. abies*) (n = 6) (right panel). First, gas exchange was measured at the soil surface, followed by two sequential measurements of root VOCs (T1, T2), each for the rhizosphere and for the ethanol group. Roots were either analyzed with their intact rhizosphere, or washed with water or 70 vol. % ethanol. Finally, the roots were removed from soil samples, sieved to 2 mm, and used for bulk soil measurements.

#### VOC measurements from the soil surface

The VOC emissions from the soil surface of all experimental plants were analyzed using open bottom chambers connected to an automated measurement system (described below). Therefore, two weeks prior to the measurements, the soil surface was cleaned of litter and moss, and the outline of the open bottom chambers (Ø 7.7 cm) was carefully trenched to a depth of 3 cm to prevent disturbance of the soil and roots on the day of measurements. The day before soil surface measurements, soil moisture was adjusted to approximately 24 vol. %.

#### Root VOC measurements

For the root VOC measurements, a part of the root system was excavated while the majority of roots remained undisturbed in the soil. For the rhizosphere group, bulk soil was carefully shaken off while preserving the rhizosphere soil. In contrast, water-washed roots were rinsed with bidistilled water until the washing solution was free of soil particles. For the ethanol group, previously water-washed roots were immersed in a 70% (v/v) ethanol solution for 30 s and subsequently rinsed with 0.5 L of bidistilled water to ensure the removal of residual ethanol.

Ethanol is among the most commonly used substances for the chemical sterilization of plant tissues, with most protocols employing concentrations of 70–80%, as this range provides an optimal balance between high disinfection efficiency and reduced dehydration compared to higher ethanol concentrations (Bodenhausen *et al*., 2013; Sahu *et al*., 2022). Compared to stronger oxidants such as sodium hypochlorite or hydrogen peroxide, ethanol represents a milder treatment with relatively minor effects on the plant tissues (Sahu *et al*., 2022). The aim of the washing treatments was to establish clear difference in the abundance of root adhering microbes across unwashed (rhizosphere), water-washed, and ethanol-washed roots, while minimizing disturbance to root tissues rather than achieving complete sterilization (Richter-Heitmann *et al*., 2016). To assess potential impacts of ethanol treatment on plant physiology, root respiration and leaf gas exchange were monitored (see description below) in in all three treatments (rhizosphere, water and ethanol-washed roots), and no treatment effect was observed on these parameters (Fig. S1).

The excavated root bundles, which remained attached to the plants, were inserted into glass cuvettes, filled with glass beads (cleaned for 2 h in an ultrasonic bath at 80°C), sealed, shaded, and connected to the automated measurement system. For a detailed description of the root enclosures, see Meischner *et al*. (2026). After two days of measurements during the first cycle (T1), the cuvettes were opened and roots were subjected to their designated washing procedures (water or ethanol washing) before measurements were resumed for the second cycle (T2).

#### VOC emissions form the bulk soil

VOC emissions and soil respiration were measured from bulk soil samples containing no living roots. To prepare these samples, soil from each of the 24 pots was sieved to 2 mm and dried at 35°C for 48 h. The soil was then transferred into glass cuvettes and adjusted to 16% gravimetric soil moisture (corresponding to 24% volumetric soil moisture at a soil density of 0.84 ± 0.01 g cm^-3^) to ensure comparability with the soil surface measurements.

Each measurement cycle lasted 24 hours (00:00 – 24:00), starting the day after open bottom chambers were installed for soil measurements on the prepared chamber outline, or two days after cuvette installation for root and shoot measurements, to minimize handling effects. Soil moisture was recorded every 30 minutes throughout the experiment (ECH2O EC-5, METER Group, Pullmann, WA, USA). After completion of the gas-exchange measurements, the dry weight and horizontally projected root and leaf/needle area was determined (image analysis software, GSA, Rostock, Germany), as well as the dry weight of absolute aboveground and belowground biomass. Soil samples were further characterized by determining their pH in water and C and N contents and isotopic ratios (δ^13^C and δ^15^N) using an elemental analyzer (Vario Isotope Cube, Elementar, Langenselbold, Germany) coupled to an isotope ratio mass spectrometer (IsoPrime, Elementar, Langenselbold, Germany) as detailed in Werner *et al*. (2009) (see Methods S1).

### Automated flow-through gas measurement system

The automated flow-through gas system for continuous measurements of VOC, CO_2_ and H_2_O fluxes was installed in two walk-in climate chambers and has been described in detail by Werner *et al*., (2020) and Meischner *et al*. (2026). It comprises a custom-built zero air generator, an analyzer unit, and 13 inlets and outlets for the connection of soil chambers or plant enclosures. A continuous air flow of 350 mL min⁻¹ was supplied to each enclosure. The incoming air was purged of VOCs via a platinum catalyst and active charcoal filter and adjusted to 430 ppm CO₂. Real-time concentrations of CO₂, H₂O, and VOCs in both the in- and outgoing air was monitored using a non-dispersive infrared gas analyzer (LI-850, LI-COR Environmental, Bad Homburg, Germany) and a proton-transfer-reaction time-of-flight mass spectrometer (PTR-TOF-MS 4000 ultra, Ionicon Analytic, Innsbruck, Austria). The PTR-TOF-MS was operated using H_3_O^+^ ionization mode. Drift tube conditions were set to a temperature of 80 °C, a pressure of 2.7 mbar, and a voltage of 503 V, resulting in an E/N of 128 Td (where E represents the electric field strength and N the number density of the buffer gas molecules in the drift tube). One empty enclosure served as a blank. A multi-position valve (VICI-Valco, Houston, TX, USA) enabled switching between the 13 system positions every 8 minutes, with a minimum instrument measurement interval of 20 seconds. The described setup allowed for the simultaneous analysis of 6 plants (2 enclosures needed per plant, one for root and one for shoot measurements) or 12 soil samples, resulting in 12 runs to measure all 4 measurement cycles for 6 replicates per species and group (rhizosphere/ethanol). All plant and soil surface measurements were conducted between September 15 – October 19, 2025 and bulk soil measurements were conducted from November 17-20, 2025.

### Data acquisition and processing

Raw data of the PTR-TOF-MS were processed with IDA software (version 2.2.0.7, Ionicon, Innsbruck, Austria) and compound assignment was performed using the GLOVOCs database (Yáñez-Serrano *et al*., 2021). The nine compounds with the highest emission rates were selected for further analysis, representing a broad spectrum of chemical classes, including hydrocarbons (e.g. mono-, and sesquiterpenes and benzene) and oxygenated compounds (e.g. oxygenated mono- and sesquiterpenes, acetone and so called green leaf volatiles). These compounds originate from diverse biosynthetic pathways, including the mevalonate (MVA) and methylerythritol phosphate (MEP) pathways (terpenes), the LOX pathway (green leaf volatiles), the shikimate pathway (benzene), and primary metabolism (acetone). Ethanol and acetaldehyde we excluded from further analysis, since their emissions are likely to be affected by the ethanol treatment.

Quantification of VOCs measured with the PTR-TOF-MS was performed using a multi-component gas mixture (Apel Riemer Environmental, USA; see Methods S3) and a Liquid Calibration Unit (LCU, Ionicon, Innsbruck, Austria). The acquired transmission data were then uploaded to the IDA software quantification module to allow for the conversion of signal intensity (counts per second) to concentrations (ppb). For further data processing and statistical analyses, the software R (version 4.2.1, R Core Team, 2021) was used. First, the average of each 8-minute measurement was calculated, excluding the first two minutes to avoid transition effects from the previous measurement. Then, the background (determined by interpolating between the blank measurements) was subtracted from root and soil measurements before proceeding with the calculations of VOC fluxes. All formulas used for the calculation of VOC emission rates (nmol m ^−2^ h^−1^) and gas exchange parameters are documented in the R-package “gasexchange3”, which is online available codeberg.org/mmEcophys/GasExchangeFormulas.git. VOC emission rates were converted to ng cm⁻² h⁻¹ using the molecular weight of each compound for consistency with GC–MS data.

### GC-MS Analysis of VOCs

In addition to continuous VOC measurements, VOC samples of roots and soils were taken for compound specific analysis of terpenoids via gas chromatography-mass spectrometry (GC-MS) (Fig. 1). For this purpose, VOCs were collected from the outlets of root enclosures or soil chambers for 150 min (09:30–12:00 h) at a flow rate of 90 mL min⁻¹ (Pocket Pump TOUCH, SKC, Dorset, UK) using glass tubes filled with 85 mg of Tenax TA (60–80 mesh; Sigma, Munich, Germany). The accumulated VOCs were subsequently analyzed via GC-MS (GC 7820A; mass-selective detector 5975, Agilent Technologies Böblingen, Germany), as described in detail by Kreuzwieser *et al*. (2025). Briefly, VOCs were desorbed at 240 °C using a thermodesorption unit and trapped at -70 °C in a cold injection system (TDU-CIS4, Gerstel, Germany). The CIS was subsequently heated to 240 °C to release the VOCs onto a DB-5MS UI 122-5532UIE column (30 m × 0.25 mm ID, 0.25 μm film thickness, Agilent Technologies, Böblingen, Germany). Helium served as the carrier gas at a constant flow of 1 mL min⁻¹. The oven temperature was increased in steps of 2 °C between 45 and 80 °C, 4 °C between 80 and 140 °C, and 9 °C between 140 and 280 °C. The mass spectrometer was operated with an ionization energy of 70 MeV, an ion source temperature of 230 °C, and a quadrupole temperature of 150 °C, recording mass spectra from 40 to 300 *m/z*. Data were processed using MassHunter software (Agilent Technologies, Böblingen, Germany). For this step, data were processed separately for spruce roots, beech roots or soil samples. Peak identification was based on the NIST library (2017) and for quantification a linear calibration curve derived from authentic standards (Sigma-Aldrich, Germany, see Methods S4) was used. For VOCs not contained in the calibration standard, standards were assigned based on chemical similarities (ring count, i.e. a-, mono- or bicyclic and functional group, i.e. pure hydrocarbons, alcohols, aldehydes, etc.), as specified in Table S1, along with match factors of each compound. VOC emission rates (ng cm ^−2^ h^−1^) were calculated based on the VOC concentrations in sampling tubes, the root or soil surface area, and the sampling duration.

### Statistical analysis

Species effects of *P. abies* and *F. sylvatica* on total VOC emissions from the soil surface and bulk soil, as well as on the compounds with the highest emission rates (α-pinene, β-pinene, β-myrcene, and α-phellandrene), were analyzed using unpaired Welch’s t-tests (n = 12) on log-transformed VOC emission rates. Log transformation was applied to account for large differences in absolute VOC emission rates between soils of both species. Species effects on soil parameters (pH, C/N ratio, and isotopic signature), which showed more homogeneous distributions across samples, were evaluated using unpaired Student’s t-tests (n = 12).

The influence of the rhizosphere on root VOC emissions was assessed by comparing rhizosphere and ethanol treatments with water-washed roots within each measurement cycle (T1 and T2). Statistical analyses were conducted within each measurement cycle to avoid confounding treatment effects with potential temporal effects between T1 and T2. Differences in mean emission rates of all compounds detected by GC–MS were analyzed between treatments and controls using unpaired Student’s t-tests with Bonferroni correction (n = 6). Assumptions of normal distribution and homogeneity of variance were assessed visually using boxplots for each compound. Statistical analyses were conducted using the “omu_summary*”* function from the R package “omu” (Tiffany & Bäumler, 2019). Log_2_fold changes and p-values are presented in a volcano plot (Fig. 4) generated with “ggplot2” (Wickham, 2016), while bar plots illustrating compounds with significant treatment effects are shown in Fig. 5. Significance levels of p < 0.05 (*), p < 0.01 (**), and p < 0.001 (***) and p < 0.1 to indicate tendencies were applied throughout.

## Results

### Soil surface and bulk soil VOC emissions associated with *P. abies* and *F. sylvatica*

For the litter-free soil surface (Fig. 1Aa), VOC emission rates and their compositions differed markedly between soils rooted by *P. abies* and *F. sylvatica* (Fig. 2A, Table 1A), with the net VOC emissions from the intact soil surface being 35-times higher in soils of *P. abies* than *F. sylvatica* (p < 0.001 ***). In *P. abies* associated soils, the mean total VOC emission rate from the soil surface was 25 ± 11 ng cm⁻² h⁻¹, of which 90% consisted of monoterpenoids and only 0.3 % of sesquiterpenoids. The dominant compounds were α-pinene, β-pinene, β-myrcene, and α-phellandrene accounting for 78 % of total VOC emissions (Fig 2A). By contrast, soils rooted by *F. sylvatica* emitted substantially lower amounts of terpenoids from the soil surface with total monoterpenoid emissions < 0.2 ng cm⁻² h⁻¹.

**Figure 2:**
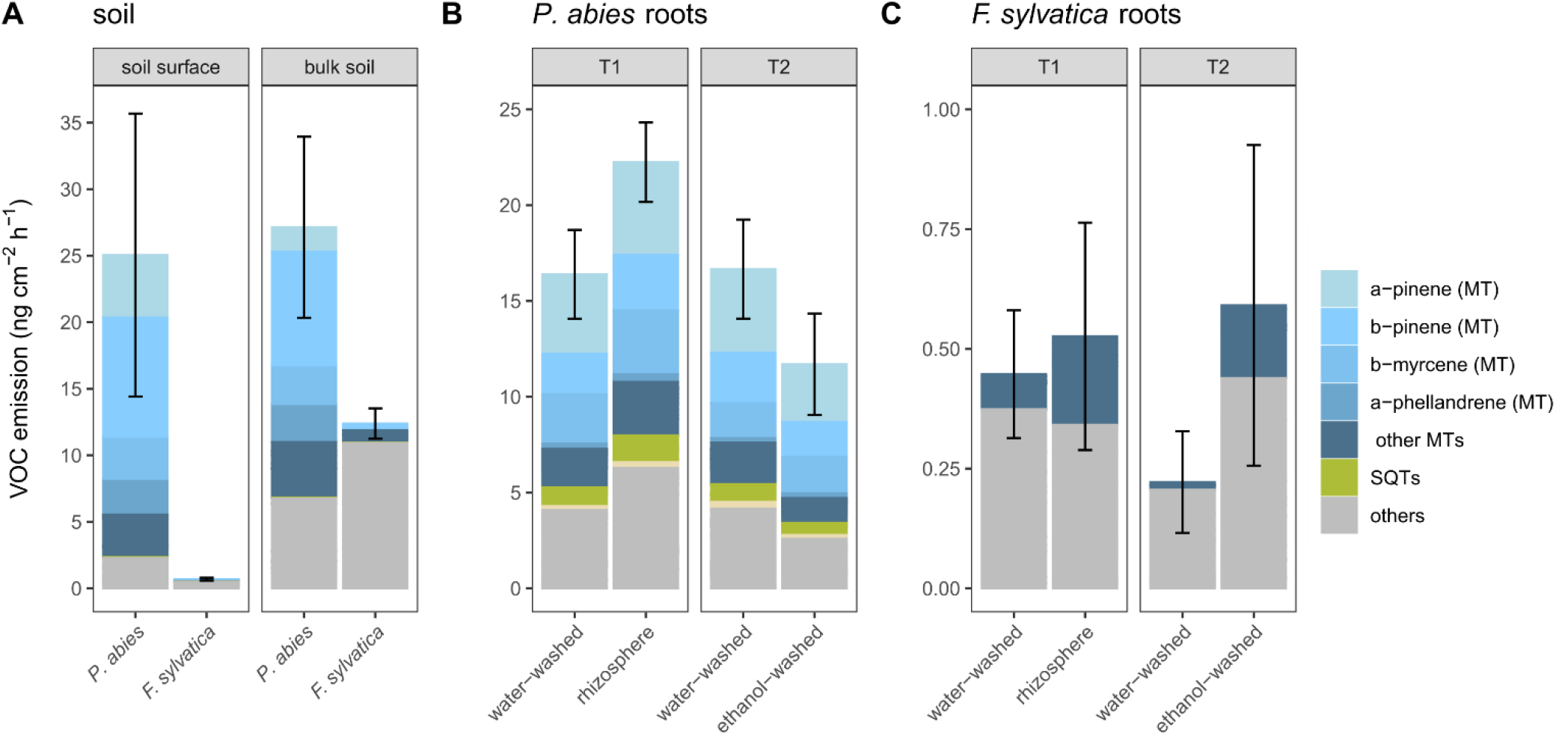
VOC emission rates from the soil surface and associated soils of *Picea abies* and *Fagus sylvatica* (A). Statistical analysis is presented in Table 1. Effects of root treatments (intact rhizosphere and ethanol washing) compared to water-washed roots (B, C). The error bars show standard deviation for total VOC emission (n = 6). T1 and T2 refer to the two experimental time periods of root VOC measurements (see Fig. 1). Note different y-axis scales in A, B and C. MT = monoterpenoids, SQT = sesquiterpenoids.

**Table 1:**
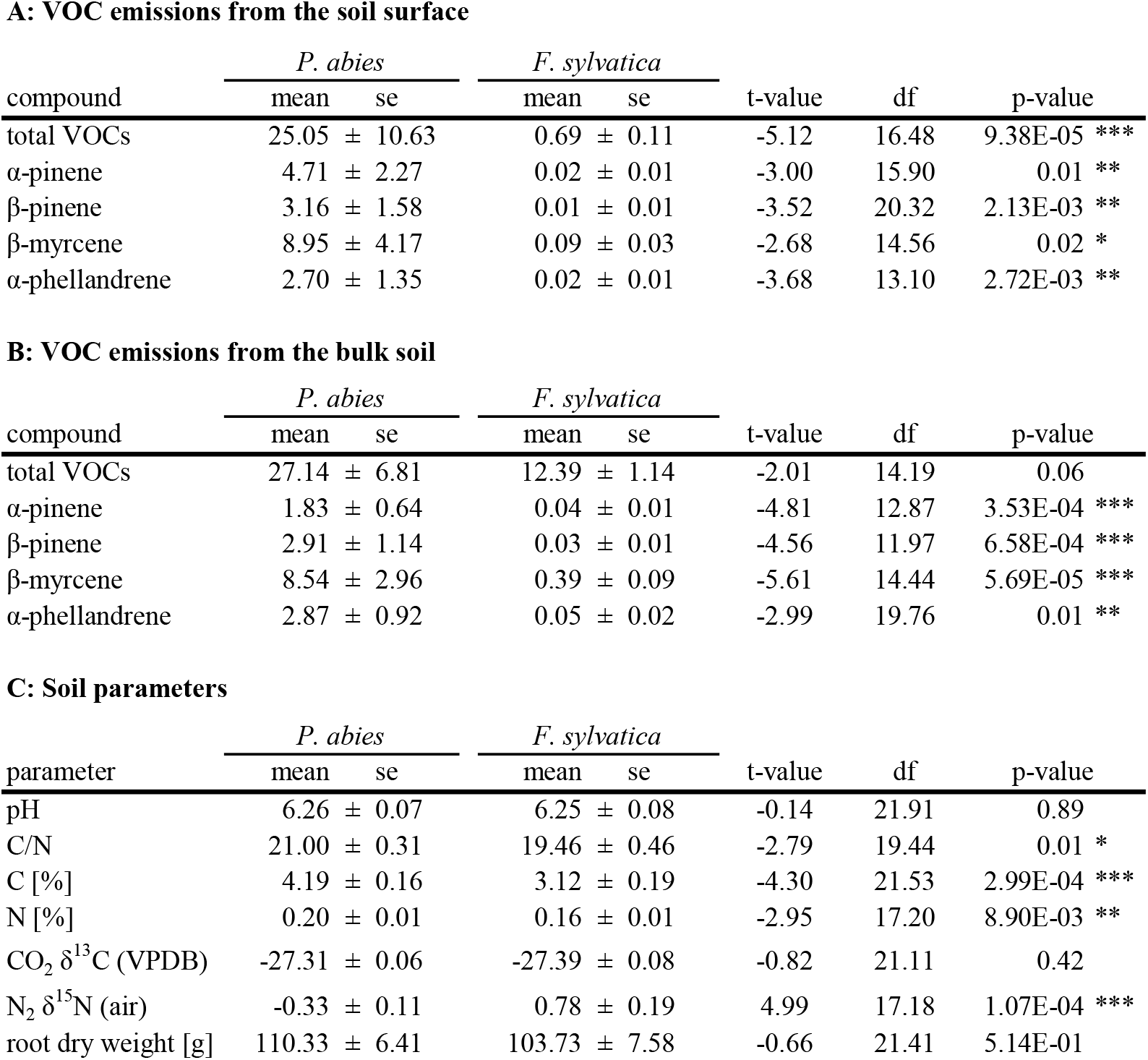
Comparison of VOC emission rates (ng cm⁻² h⁻¹) from the soil surface (A) and bulk soil (B), as well as soil parameters (C), between *Picea abies* and *Fagus sylvatica*. Four compounds with the highest overall emission rates are shown. Statistical analyses were performed on log-transformed VOC emission rates using unpaired Welch’s t-tests, and on untransformed soil parameters using unpaired Student’s t-tests. The number of replicates was n = 12 for all tests.

**A: VOC emissions from the soil surface**
| compound | *P. abies* |  | *F. sylvatica* |  | t-value | df | p-value |
| --- | --- | --- | --- | --- | --- | --- | --- |
|  | mean | se | mean | se |  |  |  |
| total VOCs | 25.05 ± 10.63 |  | 0.69 ± 0.11 |  | -5.12 | 16.48 | 9.38E-05 *** |
| α-pinene | 4.71 ± 2.27 |  | 0.02 ± 0.01 |  | -3.00 | 15.90 | 0.01 ** |
| β-pinene | 3.16 ± 1.58 |  | 0.01 ± 0.01 |  | -3.52 | 20.32 | 2.13E-03 ** |
| β-myrcene | 8.95 ± 4.17 |  | 0.09 ± 0.03 |  | -2.68 | 14.56 | 0.02 * |
| α-phellandrene | 2.70 ± 1.35 |  | 0.02 ± 0.01 |  | -3.68 | 13.10 | 2.72E-03 ** |

**B: VOC emissions from the bulk soil**
| compound | *P. abies* |  | *F. sylvatica* |  | t-value | df | p-value |
| --- | --- | --- | --- | --- | --- | --- | --- |
|  | mean | se | mean | se |  |  |  |
| total VOCs | 27.14 ± 6.81 |  | 12.39 ± 1.14 |  | -2.01 | 14.19 | 0.06 |
| α-pinene | 1.83 ± 0.64 |  | 0.04 ± 0.01 |  | -4.81 | 12.87 | 3.53E-04 *** |
| β-pinene | 2.91 ± 1.14 |  | 0.03 ± 0.01 |  | -4.56 | 11.97 | 6.58E-04 *** |
| β-myrcene | 8.54 ± 2.96 |  | 0.39 ± 0.09 |  | -5.61 | 14.44 | 5.69E-05 *** |
| α-phellandrene | 2.87 ± 0.92 |  | 0.05 ± 0.02 |  | -2.99 | 19.76 | 0.01 ** |

**C: Soil parameters**
| parameter | *P. abies* |  | *F. sylvatica* |  | t-value | df | p-value |
| --- | --- | --- | --- | --- | --- | --- | --- |
|  | mean | se | mean | se |  |  |  |
| pH | 6.26 ± 0.07 |  | 6.25 ± 0.08 |  | -0.14 | 21.91 | 0.89 |
| C/N | 21.00 ± 0.31 |  | 19.46 ± 0.46 |  | -2.79 | 19.44 | 0.01 * |
| C [%] | 4.19 ± 0.16 |  | 3.12 ± 0.19 |  | -4.30 | 21.53 | 2.99E-04 *** |
| N [%] | 0.20 ± 0.01 |  | 0.16 ± 0.01 |  | -2.95 | 17.20 | 8.90E-03 ** |
| CO<sub>2</sub> δ<sup>13</sup>C (VPDB) | -27.31 ± 0.06 |  | -27.39 ± 0.08 |  | -0.82 | 21.11 | 0.42 |
| N<sub>2</sub> δ<sup>15</sup>N (air) | -0.33 ± 0.11 |  | 0.78 ± 0.19 |  | 4.99 | 17.18 | 1.07E-04 *** |
| root dry weight [g] | 110.33 ± 6.41 |  | 103.73 ± 7.58 |  | -0.66 | 21.41 | 5.14E-01 |

The species-specific patterns persisted after removal of roots, as shown by VOC measurements on bulk soil (Fig. 2A, Table 1B). Because the handling required for bulk-soil analysis (sieving and root removal) inevitably introduces artefacts, VOC fluxes from the soil surface and bulk soil cannot be directly compared. Nevertheless, the procedure is robust for studying species effects on soil VOC emissions within each measurement approach. In the bulk soil, monoterpenoid emissions from *F. sylvatica* soils remained low, being 0.5 ± 0.1 ng cm⁻² h⁻¹ on average, and thus only a minor fraction of those from *P. abies* soils.

Although total root biomass of both species was similar (Table 1C), soil chemical properties were modulated by the tree species (Table 1C), with the. While soil pH (6.3 ± 0.2) and δ¹³C values (27.4 ± 0.2 ‰) did not differ between the two species, the soils associated with *P. abies* were enriched by 25% in carbon, resulting in a significantly higher C/N ratio of 21.0 ± 0.3 compared to 19.5 ± 0.5 in soils associated with *F. sylvatica*. Furthermore, δ¹⁵N values in *P. abies* soils were by 1.1 ‰ lower (p<0.001***) than those in *F. sylvatica* soils (Table 1C).

### Root VOC emissions under different washing treatments

Monitoring of root VOC emissions of *P. abies* and *F. sylvatica* with high temporal resolution revealed diurnal cycles in terpenoids (Fig. 2A-D), oxygenated compounds (e.g., acetone, Fig. 3H) and aromatic compounds, such as benzene (Fig. 3G). The diurnal cycles of root VOCs followed the light and air temperature regime with higher emission rates during the light phase under warmer air temperatures (25°C) than during the cooler nights (15°C) and were also apparent in the root respiration (Fig. S1C), which is a proxy for root activity.

**Figure 3:**
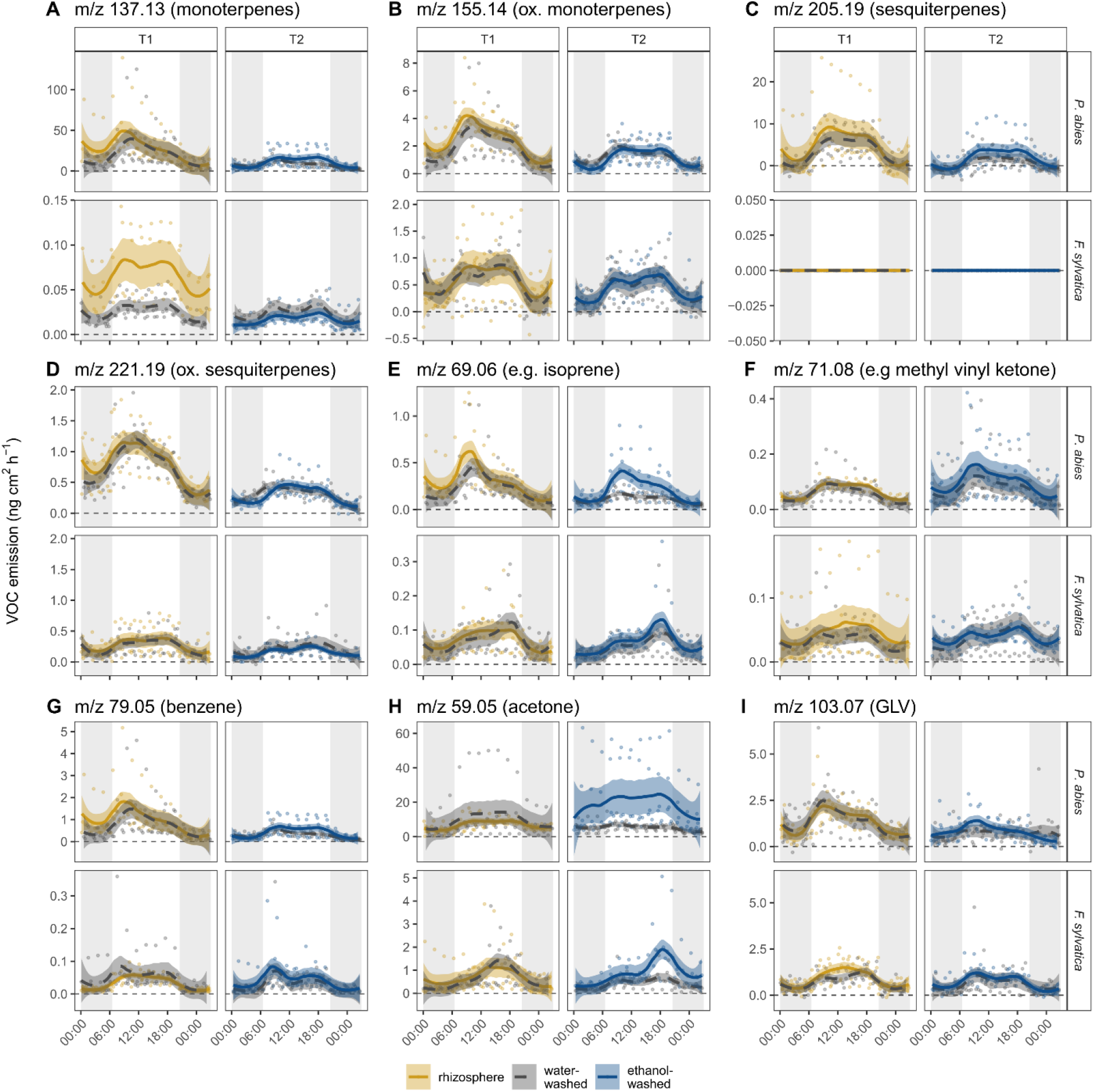
Time series of root VOC emissions from *Picea abies* and *Fagus sylvatica*. Root VOC emissions were measured from roots subjected to three root treatments (n=6 each): intact rhizosphere (yellow), water washing (gray), and ethanol washing (blue). Nonparametric local regressions (solid and dashed lines) with corresponding 95% confidence intervals were fitted using a LOESS function (smoothing parameter α = 0.4). The x-axis shows the time of day (hh:mm). Note different y-axis scales. Ox. = oxygenated. GLV = green leaf volatile. Panel titles indicate measured protonated mass to charge ratios (m/z) of VOCs and assigned compound names.

While the diurnal patterns of these VOCs were similar in *P. abies* and *F. sylvatica*, absolute daytime emission rates differed markedly, particularly for terpenoids. Daytime emissions of monoterpenes from *P. abies* roots exceeded those from *F. sylvatica* by a factor of 678, averaged across all washing treatments. Oxygenated mono- and sesquiterpene emissions from *P. abies* roots were higher by factors of 3 and 2.5, respectively, than those from *F. sylvatica* roots. Sesquiterpenes were detected to be emitted from *P. abies* roots at mean daytime emission rates of 4.7 ± 0.4 ng cm⁻² h⁻¹, but were not detectable in *F. sylvatica* roots. Similarly, emissions of other compounds such as acetone and benzene, were 16- and 15-fold higher in *P. abies*, respectively, across all washing treatments. In contrast, daytime emission rates of a green leaf volatiles (GLVs) with m/z 103.05 were comparable between the two species, with mean emission rates of 1.2 ± 0.04 ng cm⁻² h⁻¹ across both species.

From the time series of VOC emissions, it became apparent that the effect of the rhizosphere on root VOC emissions differed among compounds. Some compounds decreased, others remained constant, and some increased following removal of the rhizosphere (Fig. 3): In *F. sylvatica*, daytime monoterpene emissions from roots with an intact rhizosphere were approximately twice as high as those from water-washed roots. This indicates a net addition of monoterpenes to the VOC blend by the rhizosphere (Fig. 3A). For several root VOCs, such as benzene (Fig. 3G) and GLVs m/z 103.1 (Fig. 3I), no differences were observed between rhizosphere, water washing, and ethanol washing treatments, suggesting that these compounds passed through the rhizosphere without significant net additions or losses. In contrast, emissions of a third group of compounds increased following washing, suggesting that they were degraded within the rhizosphere. This pattern was observed for acetone in both *P. abies* and *F. sylvatica* roots. Notably, in *P. abies*, ethanol washing also led to an increase in compounds detected at m/z 69 (Fig. 3E), corresponding to the parent ion of isoprene. Mean daytime emission rates of this compounds increased 2.2-fold, from 0.14 ± 0.01 to 0.32 ± 0.03 ng cm⁻² h⁻¹, in *P. abies* roots after ethanol washing compared to water-washed roots.

Compound-specific analysis of root VOCs via GC-MS showed that a total of 59 VOCs were emitted from *P. abies* roots. Of all emitted compounds, 19 (32 %) were monoterpenoids, 7 (12 %) sesquiterpenoids (such as α-longipinene and humulene), 4 (7 %) diterpenes, with the remaining 49 % consisting of non-terpenoids or unassigned compounds (Fig. B). Diterpenes included compounds tentatively assigned as rimuene, sclarene, manoyl oxide, and isopimaradiene. Total VOC emission rates were by 33% higher in the roots with rhizosphere compared to water-washed roots and decreased by a further 33% following ethanol washing of the roots (Fig. 1B). Consistent with observations from the soil VOC analysis, monoterpenoids contributed the largest proportion of total VOC emissions from roots, accounting for 68 % across all washing treatments (Fig. 1B). The predominant compounds were again α-pinene, β-pinene, β-myrcene, and α-phellandrene (Fig. 1B).

Overall, 13 (22 %) of the detected compounds from *P. abies* roots were sensitive to rhizosphere removal (Fig. 4A, 5A). Only one compound exhibited reduced emission rates in the rhizosphere treatment relative to water-washed roots, whereas all others showed either higher emissions in the rhizosphere compared to water-washed roots or higher emissions in the water-washed roots compared to ethanol-washed roots (Fig. 4A). For some compounds, such as α-terpinolene, emissions decreased stepwise with increasing washing intensity: levels were by 23 % lower in water-washed roots than in roots with intact rhizosphere (from 0.21 ± 0.2 to 0.16 ± 0.8 ng cm⁻² h⁻¹, p = 0.006 **) and further decreased by 55 % (from 0.09 ± 0.03 to 0.04 ± 0.02 ng cm⁻² h⁻¹, p = 0.009 **) following ethanol washing (Fig. 4A, Fig. 5A). Notably, β-myrcene was the compound with the highest emission rate in *P. abies* roots that declined after water washing, decreasing from 3.33 ± 0.6 to 2.56 ± 0.6 ng cm⁻² h⁻¹, p = 0.048 *). Thus, emission rates of VOCs, in particular monoterpenoids, from *P. abies* roots generally decreased with the degree of removal of the rhizosphere.

**Figure 4:**
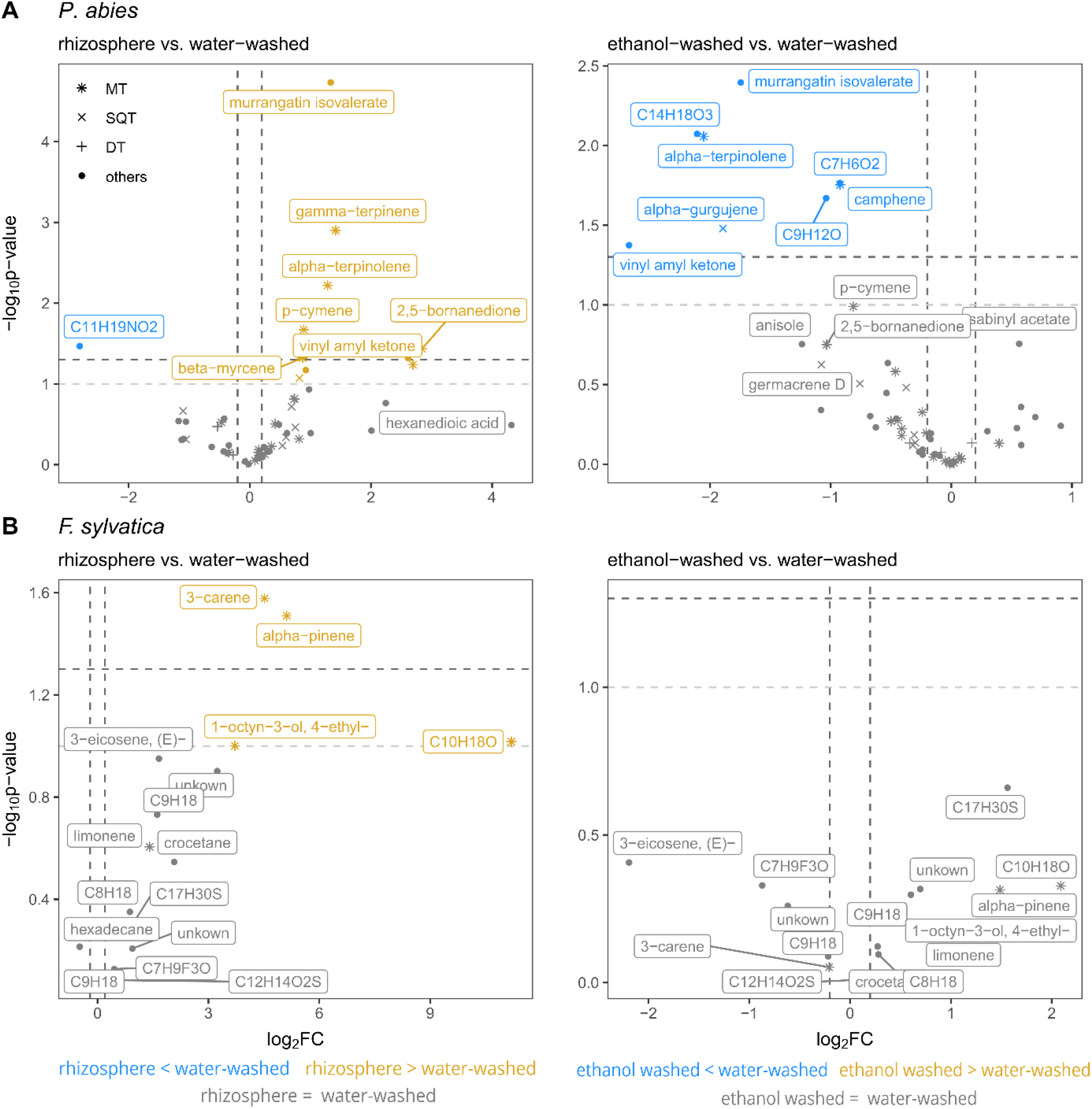
Volcano plots of all detected VOCs emitted from *Picea abies* (panel A) and *Fagus sylvatica* (panel B) roots. Results of unpaired Student’s t-tests (n = 6) comparing roots with intact rhizosphere vs. water-washed roots (left panels) and ethanol-washed roots vs. water-washed roots (right panels) are shown. The y-axis indicates the significance level (– log₁₀-transformed p-value), and the x-axis represents the effect size (log₂ fold change) of differences between the compared treatments. Horizontal dotted lines denote significance thresholds (dark gray: p = 0.05, significant; light gray: p < 0.1, indicating a tendency), whereas vertical dotted lines mark log₂ fold change thresholds (−0.2 and 0.2). Compounds meeting both the tendency (p < 0.1) and effect size criteria are highlighted, with positive log₂ fold changes (increased emissions relative to water-washed roots) shown in orange and negative changes (decreased emissions relative to water-washed roots) in blue. MT = monoterpenoids, SQT = sesquiterpenoids, DT = diterpenoids, others = non-terpenoids and unassigned compounds.

**Figure 5:**
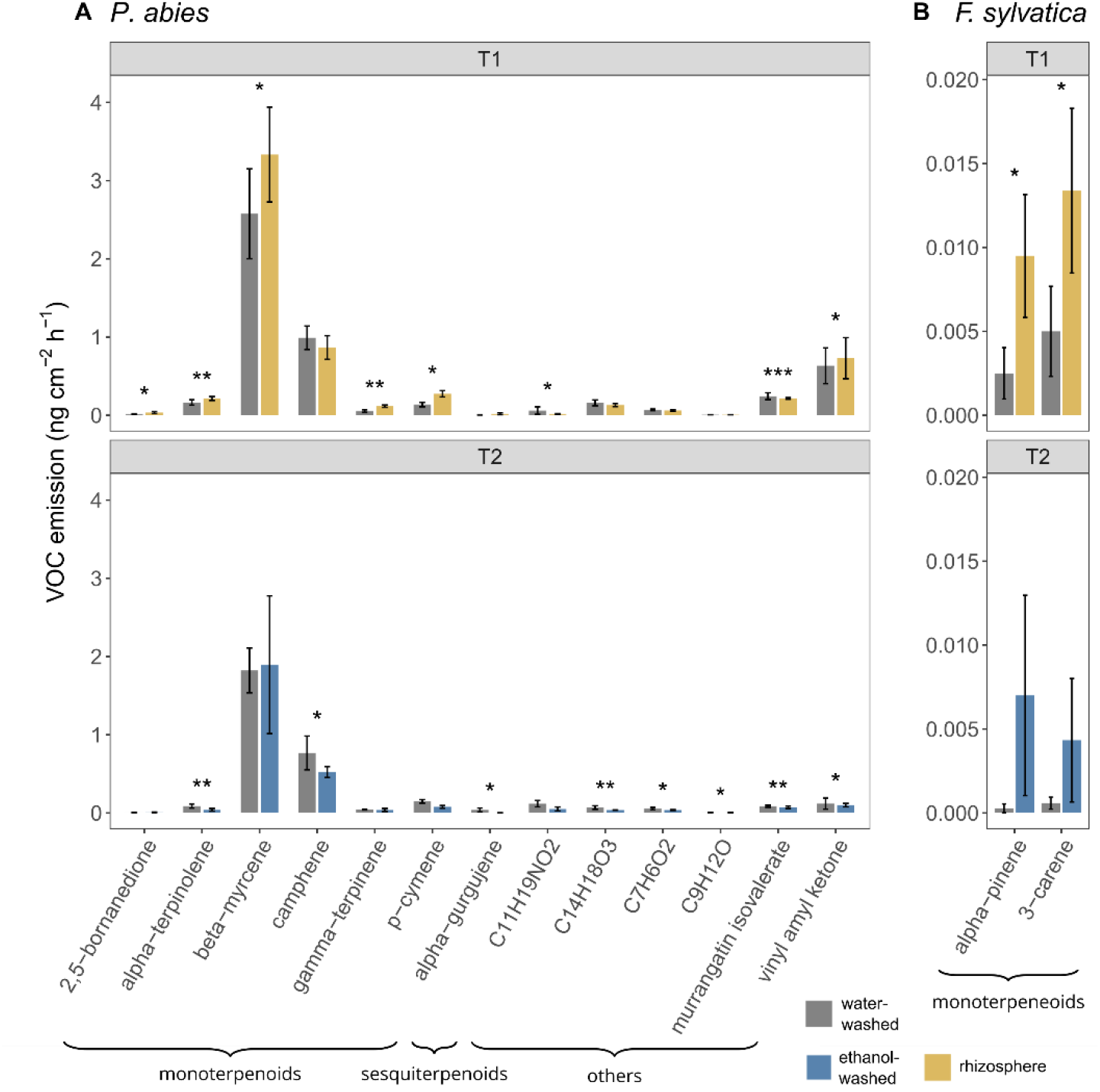
Bar plots of root VOC emissions from *Picea abies* (panel A) and *Fagus sylvatica* (panel B) that were sensitive to root washing treatments. Asterisks indicate significant differences between treatments (rhizosphere vs. water-washed or ethanol-washed vs. water-washed). Statistical analyses were performed using unpaired Student’s t-tests. Significance levels are indicated as p < 0.05 (*), p < 0.01 (), and p < 0.001 (*). T1 and T2 refer to the two experimental time periods of root VOC measurements (see Fig. 1).

A similar, though less pronounced, pattern was observed in *F. sylvatica* roots. Of the 16 compounds detected, with overall low total emission rates ranging between 0.2 and 0.5 ng cm⁻² h⁻¹, three (19 %) were monoterpenes, namely α-pinene, 3-carene and limonene. Emissions of α-pinene and 3-carene were significantly lower in the water-washed roots compared to roots with intact rhizosphere (3-carene, p = 0.03 *; α-pinene, p = 0.03 *), and two additional non-terpenoids showed a tendency toward lower emissions in the water-washed roots compared to the rhizosphere treatment (p < 0.1). No further decreases in emissions of any detected compounds were observed following additional ethanol washing.

In summary, these results indicate that VOC production, particularly of monoterpenoids, exceeded their consumption in the rhizosphere. While the majority of root-emitted VOCs passed through the rhizosphere without significant loss or addition, a distinct pattern emerged for monoterpenoids, whose emissions were consistently enhanced in rhizosphere soil. This suggests that the rhizosphere acts as a hotspot of monoterpenoid production rather than a sink. In contrast, only a few compounds, potentially including isoprene, decreased by passing the rhizosphere.

## Discussion

We investigated VOC emissions at the root–soil and soil–atmosphere interfaces, with a focus on the rhizosphere of two temperate tree species. Species-specific emission patterns were observed in roots and soils, characterized by a diverse, terpenoid-rich VOC composition with higher total emission rates in *P. abies*, whereas *F. sylvatica* exhibited a less diverse VOC profile with lower emission rates and contribution of terpenoids (Fig. 2, Fig). These patterns were evident not only in root emissions but also, notably, at the soil surface and in root-free bulk soil (Fig. 2). By isolating bulk soil, root plus rhizosphere, and root emissions, we identified the rhizosphere as a hotspot of VOC production in *P. abies*, providing new insights into the sources and controls of belowground VOC emissions.

### Rhizosphere is a hotspot VOC production

The present study provides evidence that the rhizosphere acts as a source rather than a sink of terpenoids and can substantially influence the net VOC emissions measured from roots of *P. abies*, a representative for terpenoid-rich tree species. Effects of rhizosphere removal were less pronounced in *F. sylvatica* (Fig. 4, 5), a species characterized by generally lower terpenoid emission rates. This contrasts our initial hypothesis that high microbial activity in the rhizosphere would lead to a net decrease in root VOCs due to microbial degradation, but is in line with high microbial terpenoid emissions reviewed by Avalos *et al*. (2022) and reported e.g. from soil microorganisms (Jüttner, 1990) and fungal cultures (Guo *et al*., 2020). The profound effects of rhizosphere soil and its microbial communities on root VOC emissions observed in this study raise the question for future research: How to include rhizosphere effects in standardized ways in studies on root VOC emissions? Given that VOC production in the rhizosphere exceeded uptake, subjecting roots to washing procedures may lead to an underestimation of net root VOC emissions. Unfortunately, our data allow no conclusion on whether rhizosphere microorganisms directly enhanced net VOC emissions by their own VOC production, or whether they stimulated VOC production from root-tissues. For aboveground measurements, leaf VOC emissions are typically quantified with the enclosure technique without removing the phyllosphere microorganisms (Niinemets *et al*., 2011; Frey *et al*., 2025). This approach has several advantages: 1) it avoids disturbance of plant tissues through chemical or mechanical (e.g. sonication) sterilization procedures; 2) measurements are straightforward and do not require extensive sample preparation; 3) net VOC fluxes are used for integration of leaf-level VOC emissions into land surface models, such as LPJ-GUESS (Belda *et al*., 2022) and 4) removing phyllosphere microorganisms may change VOC production by plant tissues (Junker & Tholl, 2013; Bringel & Couée, 2015; Farré-Armengol *et al*., 2016). For example, *Fusarium sp*. and *Melampsora spp.* have been shown to affect VOC production by inducing an immune response by the plant (Toome *et al*., 2010; Wenda-Piesik *et al*., 2010). Similar considerations apply to studies of root VOC emissions, although the rhizosphere differs considerably from the phyllosphere as a habitat for microorganisms in terms of nutrient availability, moisture, oxygen availability, among other factors (Junker & Tholl, 2013). Numerous studies have documented close functional interactions between roots and rhizosphere microorganisms including plant-growth-promoting rhizobacteria (Berg & Smalla, 2009; Berendsen *et al*., 2012) and mycorrhizal fungi (Taylor *et al*., 2000). Mycorrhizal associations, such as between *P. abies* and ectomycorrhizal fungus *Paxillus involutus* (Marschner & Godbold, 1995; Jentschke *et al*., 2000), illustrate tight linkage of plant roots and rhizosphere microorganisms and the challenges to define a clear boundary between root tissue and its surrounding microbial community in experimental setups. Richter-Heitmann *et al*. (2016) demonstrated that even vigorous washing procedures removed only 45 % of rhizosphere microorganisms, indicating that completely sterile roots are difficult, if not impossible, to obtain under realistic experimental conditions. Moreover, although ethanol washing did not lead to any detectable effects on plant physiology in this experiment (see Fig. S1), root washing is substantially more invasive than carefully shaking off loose soil. The benefits of root washing procedures may thus only outweigh limitations when mechanistic questions on emission and degradation processes are addressed. Conversely, quantifying VOC emissions originating exclusively from the rhizosphere in the absence of roots remains methodologically even more challenging than removing microorganisms from the root surface, since many rhizosphere microorganisms are not cultivable in the laboratory (Singh *et al*., 2004).

Preserving an intact rhizosphere for root VOC measurements, might, on the other hand, lead to an underestimation of compounds that get rapidly degraded by soil microbes, such as isoprene (Dawson *et al*., 2023; Pugliese *et al*., 2023, 2026). Bacterial metabolism of isoprene is estimated to be responsible for the uptake of approximately 20.4 Tg yr^-1^ atmospheric isoprene (Cleveland & Yavitt, 1997). When isoprene production is balanced by microbial uptake, isoprene emissions become detectable only after microbial degradation is inhibited prior to VOC sampling, as has been recently shown for *Sphagnum* moss (Crombie *et al*., 2025). A similar pattern was observed in the present study: In *P. abies* roots, ethanol washing increased the abundance of a compound detected at m/z 69 and tentatively assigned as isoprene (Fig. 3E), suggesting that microbial degradation may have reduced its concentration in unwashed and water-washed roots. However, the identity of this compound could not be confirmed by GC–MS because the adsorbent material used for VOC collection in this study (Tenax-TA) does not efficiently trap isoprene (Cao & Nicholas Hewitt, 1993). Therefore, these findings should be regarded as a first indication of simultaneous isoprene production and microbial degradation in *P. abies* roots. Future experiments subjecting the roots to specific isoprene monooxygenase inhibitors, such as acetylene and 1-octyne (Dawson *et al*., 2020; Wright *et al*., 2020; Sims *et al*., 2022), combined with VOC sampling on adsorbent material with high adsorbent capacity for isoprene, such as carbotrap (Cao & Nicholas Hewitt, 1993), will help to quantify root VOC emissions and microbial consumption.

### Soil VOC emissions exhibit species-specific VOC profiles

The higher terpenoid emissions from the soil surface of *P. abies* compared to *F. sylvatica* observed in this study may be attributed to a combination of higher release rates from roots (Asensio *et al*., 2008; Wenke *et al*., 2010; Peñuelas *et al*., 2014), greater microbial (especially fungal) terpenoid production (Stahl & Parkin, 1996; Wenke *et al*., 2010; Schmidt *et al*., 2015), and enhanced release from terpenoid storage pools within the soil (Isidorov *et al*., 2010; Mäki *et al*., 2019b; Kreuzwieser *et al*., 2025). Differences in total belowground biomass is unlikely to explain this pattern, as this was similar between *P. abies* and *F. sylvatica*. Terpenoid storage pools are formed by organic matter in the upper soil, including decaying leaves and needles, root litter, and resin, as well as from the sorption of terpenoids to soil particles (Ruiz *et al*., 1998; Aochi & Farmer, 2005; Serrano & Gallego, 2006). As such, they might explain, how species-effects in terpenoid emissions persisted in the bulk soil even after removal of living roots and leaf/needle litter. (Fig. 2B). A predominantly plant-derived origin of terpenoid emissions from bulk soil, rather than primarily microbial production, is supported by the substantial overlap in compound composition and relative abundances of the dominant terpenoids, α-pinene, β-pinene, β-myrcene, and α-phellandrene, across emissions from the soil surface, bulk soil, and roots of *P. abies*. Our observation of species-specific soil VOC patterns is supported by Lee *et al*. (2025), who reported higher terpenoid emissions and greater terpenoid storage in soils beneath the conifer *Pseudotsuga menziesii* than beneath *F. sylvatica* in a forest stand (Werner *et al*., 2024; Tesch *et al*., 2025) close to our study site.

Additionally, higher soil C contents and lower δ¹⁵N signatures in *P. abies* soils compared to *F. sylvatica* soils observed in this study, demonstrate that tree species shape carbon and nitrogen turnover processes. Species-specific effects on soil biochemical processes may result directly from contrasting nutrient acquisition strategies (Augusto *et al*., 2002; Vesterdal *et al*., 2012; Prescott & Vesterdal, 2013), or indirectly from tree-mediated shifts in soil microbial community composition and function (Dukunde *et al*., 2019; Singavarapu *et al*., 2022). Also, plant derived VOCs may influence carbon and nitrogen cycling, adding another layer to the complex tree–soil interactions. For example, fumigation of soils under a *Betula pendula* stand with VOCs from *P. abies* and *Pinus sylvestris* resin increased C mineralization and decreased net N mineralization in both N-rich and N-poor forest soils (Uusitalo *et al*., 2008). The mechanisms have not yet been fully elucidated, however, they likely involve both the antimicrobial properties of terpenoids (Gershenzon & Dudareva, 2007), which can alter microbial activity and decomposition processes, and their utilization as carbon and energy sources by specialized microorganisms (Kleinheinz *et al*., 1999). Taken together, the species-specific differences in soil VOC emissions observed in this study may influence the composition and activity of soil microbial communities and *vice versa*. The processes in the rhizosphere, as the interface between roots and soil microorganisms, is therefore of particular relevance for enhancing our understanding of belowground VOC dynamics and carbon turnover.

Overall, this study demonstrates that tree-species effects on soil VOC emissions persist even after the removal of litter and living roots, as evidenced by the high and diverse terpenoid emissions from *P. abies* soils and the lower and less diverse terpenoid emissions from *F. sylvatica* soils. Furthermore, we show that the rhizosphere acts as a hotspot of terpenoid production in the coniferous species *P. abies*. While research on root exudation has largely focused on non-volatile compounds, this study contributes to extend our understanding of root exudates to include volatile compounds. Our findings further illustrate that tree species influence ecosystem VOC fluxes not only through aboveground emissions, but also through root emissions and potentially trough modified terpenoid storage pools. Notably, the compounds with the highest root and soil emission rates, α-pinene, β-pinene, β-myrcene, and α-phellandrene, are among the dominant VOCs emitted from Northern Hemisphere forest ecosystems (Geron *et al*., 2000). Consequently, observed species-specific differences in soil emissions of these compounds may affect the composition and magnitude of ecosystem-scale VOC emissions with cascading effects on the ecosystem: Terpenoids differ in their reaction rates with O₃, OH, and NO₃ radicals (Hatakeyama *et al*., 1989; Hallquist *et al*., 1999), which determine their atmospheric lifetimes and capacities to form secondary organic aerosols (Hoffmann *et al*., 1996; Atkinson, 2007). By influencing the abundance and composition of reactive terpenoids released from forest soils, tree species may therefore indirectly affect atmospheric oxidation processes, aerosol formation, and biosphere-atmosphere feedbacks. A comprehensive understanding of the production, transformation, storage, and uptake of root-derived VOCs is ultimately required to incorporate these belowground processes into process-based ecosystem models.

## Acknowledgements

The authors would like to thank Eva Schottmüller and Madita Vogl for gardening the experimental plants, Michel Grün for his assistance with data collection and Robin Schneider for performing the GC-MS analysis.

**The following supporting information is available for this article:**

Methods S1: Analysis of soil pH in water

Methods S2: Analysis of soil, C/N and δ13C and δ15N

Methods S3: Multi-component gas mixture for calibration of PTR-TOF-MS data

Methods S4: Overview of standards used for calibration of GC-MS data

Figure S1: Time series of shoot gas exchange and VOC emissions and root respiration

Figure S2: Time series of root ethanol, acetaldehyde and GLVs emissions

Table S1: Compound list of GC-MS analysis with assigned calibration standards

## Funding

German research foundation DFG (ECOSENSE – SFB 1537, project ID 459819582; Future Forest – EXC 3127, project ID 533786343)

Danish national research foundation (VOLT - Center for Volatile Interactions, project ID DNRF168)

## Author Contributions

MM designed the study with input from CW and RR. CW and RR acquired the funding for this study. MM and AS conducted the measurements and analyzed the data. MM wrote the manuscript with input from all authors.

## Data availability

The datasets generated during this study have been deposited in the Zenodo repository (DOI: 10.5281/zenodo.21888979). Access to the dataset is currently restricted for reviewers during the peer-review process. The dataset will be released publicly upon acceptance of the manuscript.

