## Supplementary material for "The rhizosphere of *Picea abies* is a hotspot of terpenoid production": Methods S1

**The following supporting information is available for this article:**

Methods S1: Analysis of soil pH in water
Methods S2: Analysis of soil, C/N and $\delta^{13}C$ and $\delta^{15}N$
Methods S3: Multi-component gas mixture for calibration of PTR-TOF-MS data
Methods S4: Overview of standards used for calibration of GC-MS data
Figure S1: Time series of shoot gas exchange and VOC emissions and root respiration
Figure S2: Time series of root ethanol, acetaldehyde and GLVs emissions
Table S1: Compound list of GC-MS analysis with assigned calibration standards

Methods S1: Analysis of soil pH in water

Soil pH was determined by preparing a 1:2.5 (w/v) suspension of soil (< 2mm, 5 g) and demineralized water, shaking and allowing the suspension to equilibrate for 4 h. Afterwards, the pH of the aqueous supernatant was measured using a microprocessor pH Meter (pH 526, WTW, Vienna, Austria) following a 2-point calibration. The electrode was rinsed with demineralized water between measurements, and temperature compensation was applied to all readings.

Methods S2: Analysis of soil, C/N and $\delta^{13}C$ and $\delta^{15}N$

For $\delta^{13}C$ and $\delta^{15}N$ analysis, samples were dried for 48 h at 60°C, pulverized and 20 mg of soil powder was transferred into tin capsules. Samples were analyzed using an elemental analyzer (EA) (Vario Isotope Cube, Elementar, Langenselbold, Germany) coupled to an isotope ratio mass spectrometer (IRMS) (IsoPrime, Elementar, Langenselbold, Germany) as detailed in Werner *et al.* (2009). EA-IRMS data were referenced to IAEA-600 (caffeine) ($\delta^{13}C$ = –27.8 ± 0.07) and $\delta^{15}N$ = 1.0 ± 0.16). The isotope ratios of the samples are expressed against international standards as δ (‰) notation according to equation (1):

$$\delta\ [‰] = \left(\frac{R_{sample}}{R_{standard}} - 1\right) * 1000 \qquad (1)$$

where $R_{sample}$ and $R_{standard}$ are isotopic ratios of a sample and the international standards (Vienna-Pee Dee Belemnite for carbon isotopes and atmospheric $N_2$ for nitrogen isotopes), respectively.

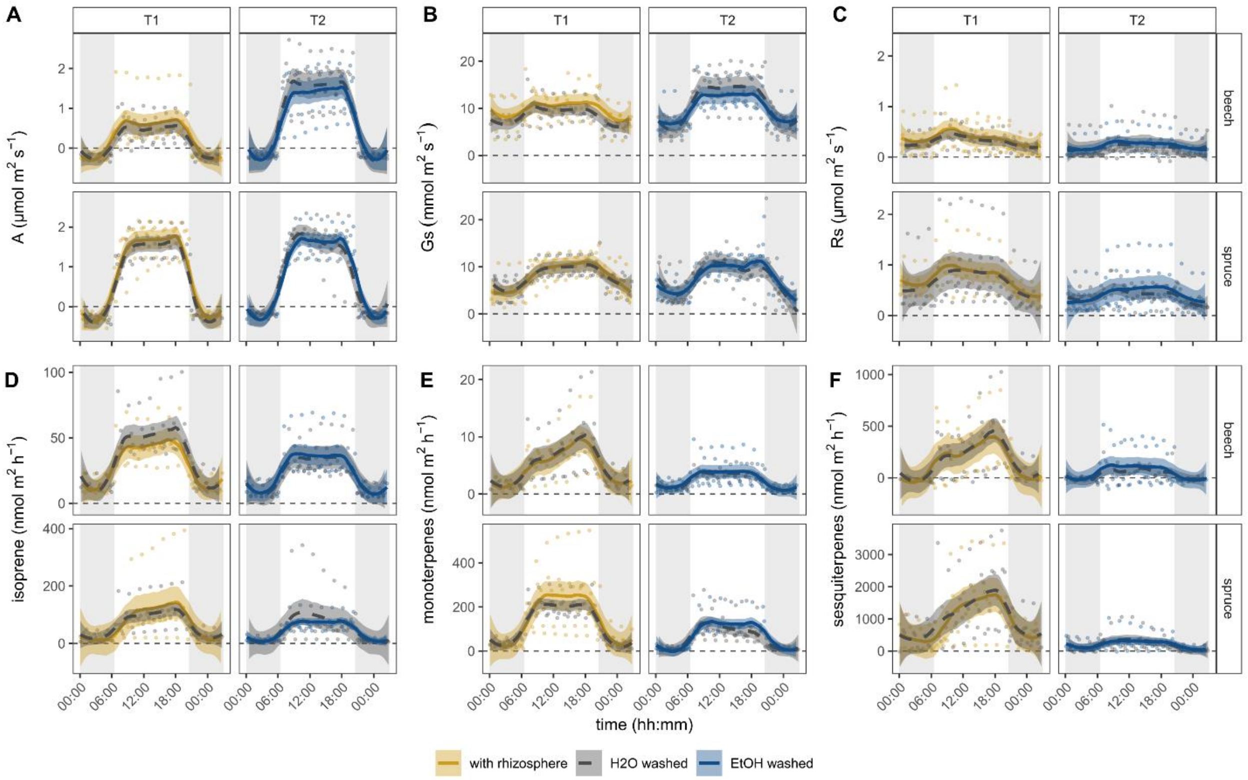


Figure S1: Time series of gas exchange from shoots (A &B) and roots (C), as well as VOC emissions from shoots of Picea abies (spruce) and Fagus sylvatica (beech) under different washing treatments (n = 6). Nonparametric local regressions (solid and dashed lines) with corresponding 95% confidence intervals were fitted using a LOESS function (smoothing parameter α = 0.4). Water-washed roots served as the control.

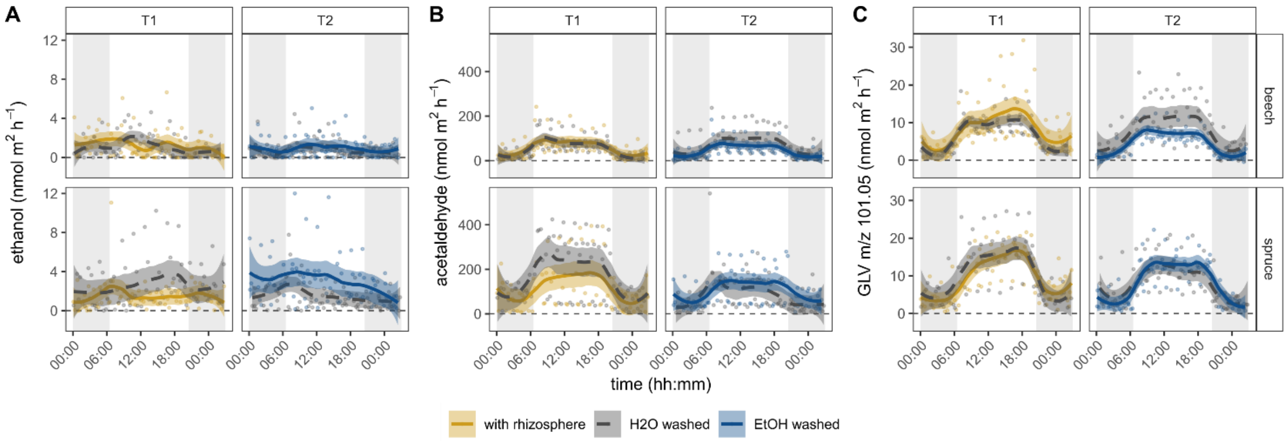


Figure S2: Time series of root VOC emissions from Picea abies (spruce) and Fagus sylvatica (beech) under different washing treatments (n = 6). Data were recorded using PTR-TOF-MS under controlled conditions. Nonparametric local regressions (solid and dashed lines) with corresponding 95% confidence intervals were fitted using a LOESS function (smoothing parameter α = 0.4). Water-washed roots served as the control.
